# Self-growth suppression in *Bradyrhizobium diazoefficiens* is caused by a diffusible antagonist

**DOI:** 10.1101/2024.06.01.596975

**Authors:** Armaan Kaur Sandhu, Brady R. Fischer, Senthil Subramanian, Adam D. Hoppe, Volker S. Brözel

## Abstract

Microbes in soil navigate interactions by recognizing kin, forming social groups, exhibiting antagonistic behavior, and engaging in competitive kin rivalry. Here, we investigated a novel phenomenon of self-growth suppression (sibling rivalry) observed in *Bradyrhizobium diazoefficiens* USDA 110. Swimming colonies of USDA 110 developed a distinct demarcation line and inter-colony zone when inoculated adjacent to each other. In addition to self, USDA 110 suppressed growth of other *Bradyrhizobium* strains and several other soil bacteria. We demonstrated that the phenomenon of sibling rivalry is due to growth suppression but not cell death. The cells in the inter-colony zone were culturable but have reduced respiratory activity, ATP levels and motility. The observed growth suppression was due to the presence of a diffusible effector compound. This effector was labile, preventing extraction, and identification, but it is unlikely a protein or a strong acid or base. This counterintuitive phenomenon of self-growth suppression suggests a strategic adaptation for conserving energy and resources in competitive soil environments. *Bradyrhizobium’s* utilization of antagonism including self-growth suppression likely provides a competitive advantage for long-term success in soil ecosystems.

## Introduction

Microbes in soil encounter numerous challenges that influence their population dynamics and success. Sporadic or seasonal nutrient input and the generally high microbial densities poses a challenge for microbial survival. Soil microbial populations displaying long-term success are, therefore, expected to use these finite resources sparingly [1]. Nutrient availability determines growth yield, and low nutrient conditions select oligotrophs over copiotrophs [2]. Spatial constraints within microbial communities are an additional challenge, limiting physical space, growth, and dispersal [3]. Microorganisms express diverse social behaviors, and many can recognize their neighboring microbial populations for cooperation or competition [4]. Microbes also face challenges from surrounding microbial communities, and many microbial taxa have evolved mechanisms to suppress their competitors [5]. Some taxa use long-range strategies to inhibit others while others use contact-dependent or short-range mechanisms. One prominent long-range example is the production of antimicrobial compounds like antibiotics and bacteriocins. Diffusion through the soil inhibits the growth of nearby competitors who then no longer consume resources, giving the antagonist a competitive advantage [6; 7; 8; 9]. Other long-range mechanisms include iron-scavenging systems to deprive competitors, suppressing their growth [10]. Examples of short range or contact dependent mechanisms include the type VI secretion system (T6SS), in syringe-like protrusions inject toxins into the competitors [6; 11; 12; 13].

Successful competitive strategies typically involves identification and differentiation between genetically related (self) and unrelated individuals (non-self) [14; 15]. Microbes have various ways of differentiating among self and non-self. The most widely researched is Quorum Sensing (QS), where population density is measured through extracellular concentration of autoinducer [16]. Some autoinducer systems are strain or species-specific, while others act across species. In biofilm co-cultures *Agrobacterium tumefaciens* biomass decreased when occurring with wild-type *Pseudomonas aeruginosa,* but remained constant with a mutant lacking QS abilities, indicating competitive interactions [17]. Another phenomenon to differentiate self and non-self is the toxin-antitoxin (TA) system, comprised of two elements: a stable toxin that inhibits a cellular process and an antitoxin of short half-life that counteracts the cognate toxin [18]. Many bacterial populations use TA systems for cell contact-dependent killing of competing species, including *Vibrio cholerae, P. aeruginosa,* and *Caulobacter crescentus* [13; 19; 20]. *Myxococcus xanthus,* a model organism for exploring social behavior, expresses SitA toxin cassettes to discriminate strains and contribute to population structure [21]. In *M. xanthus* strains that express different *sitAI* cassettes, demarcation zones form between swarming colonies [21]. Colonies originating from the same ancestral isogenic strain merge seamlessly, but a demarcation boundary becomes evident when one colony is from the ancestral isolate and the other originated a few cycles later from the same ancestor [22]. Such development of a demarcation line between nonidentical strains or species is termed kin rivalry and has been reported in swarming *Proteus mirabilis* and *Bacillus subtilis* [22; 23; 24]*. B. subtilis* was capable of forming swarming colony boundaries with less phylogenetically related soil isolates, and the isolates that exhibited boundary formation were also reported to compete for root surface colonization [23]. While kin rivalry between nonidentical strains is well established, isogenic strains of swimming *Marinobacter algicola,* and *Pseudomonas putida* and swarming *Paenibacillus dendritiformis* have been reported to develop a boundary between two colonies, termed sibling rivalry [25; 26; 27; 28]. Recently, Kastrat and Cheng reported inhibition within and across *Escherichia coli* strains and other bacteria [28]. This broad-spectrum inhibition appeared dependent on a secreted inhibitory compound as it was independent of nutrient depletion and QS, but the nature of the compound was not reported. In *P. dendritiformis*, subtilisin, a lethal protein that regulates colony density, is secreted at higher cell density, resulting in the development of a demarcation line between the two swarming colonies [29]. *M. algicola* arrest in swimming populations has been linked to the protein glycerophosphoryl diester phosphodiesterase (GDPD), but its role in this process remains unclear [25]. No chemical compounds were detected in the case of *P. putida,* where compression waves among opposing colonies cause the development of the demarcation line [27].

We unexpectedly discovered sibling rivalry between two swimming isogenic colonies of *Bradyrhizobium diazoefficiens* USDA 110, a biological nitrogen fixer in soybean root nodules [30]. While studying competition among swimming *Bradyrhizobium* strains, two isogenic colonies of USDA110 used as the negative control also paused swimming upon approaching each other, forming a demarcation line. The phenomenon of suppressing of self is counter intuitive as *Bradyrhizobium* are slow growing, and to nodulate, it must compete in a rhizosphere teaming with faster-growing taxa supported by root exudates [31]. Here, we report a) characteristics of this phenomenon and b) experiments to understand the underlying mechanism. This behavior of *B. diazoefficiens* is intriguing and unique because it includes restraining population growth of self, other *Bradyrhizobium,* and other soil bacteria. This phenomenon suggests a finely tuned strategy for limiting resource utilization as opposed to the concept of the copiotroph, the voracious species that eats as much as it can to grow its population as large as it can. These findings help explain how the slow-growing *Bradyrhizobium* is the most predominant bacterial genus in soils across the planet [32].

## Materials and Methods

### Bacterial strains and culture conditions

*Bradyrihizobium* strains, mutants of *B. diazoefficiens* USDA110, and other bacteria used are listed in Table S1. USDA110 was genetically tagged with bjGFP, mTq2, sYFP2, and mChe following the originators’ method [33]. bjGFP has been codon frequency optimized for *B. diazoefficiens* [33]. The plasmid pRJPaph-gfp, generously provided by Hans-Martin Fischer, was transformed into *E. coli* S17-1 λpir, and then subsequently transferred to USDA 110 through a biparental conjugation. USDA 110 was cultivated in PSY medium supplemented with L-(+)-arabinose (1 g/l) (Thermo Scientific A11921) [34]. Cultures were incubated at 28° C and shaken at 250 rpm. The medium was supplemented with 0.35% agar to conduct motility experiments. For all the following motility experiments, a 48 h exponential phase culture grown in PSY was used as an inoculum. We also used filter-sterilized soil extract (SESOM) as a motility medium to replicate the nutritional conditions found in soybean field soil [35; 36]. To visualize swimming, 20 µl of exponentially grown USDA 110 diluted to an absorbance (600 nm) of 0.100 were spotted onto low percentage agar plates (0.35%) and the plates were incubated for 10 d or as specified. Plates were inoculated with either 2, 4, or 16 droplets of the inoculum depending on the experiment, (Fig. 1A-C). However, for most of the subsequent experiments, 4 droplets of the inoculum were used as standard, with 2.5 cm vertical distance and 1.5 cm horizontal distance (Fig. 1E). All other strains of *Bradyrhizobium* were inoculated in the same manner. The plates were incubated at 28° C for 10 d unless stated otherwise.

**Figure 1.**
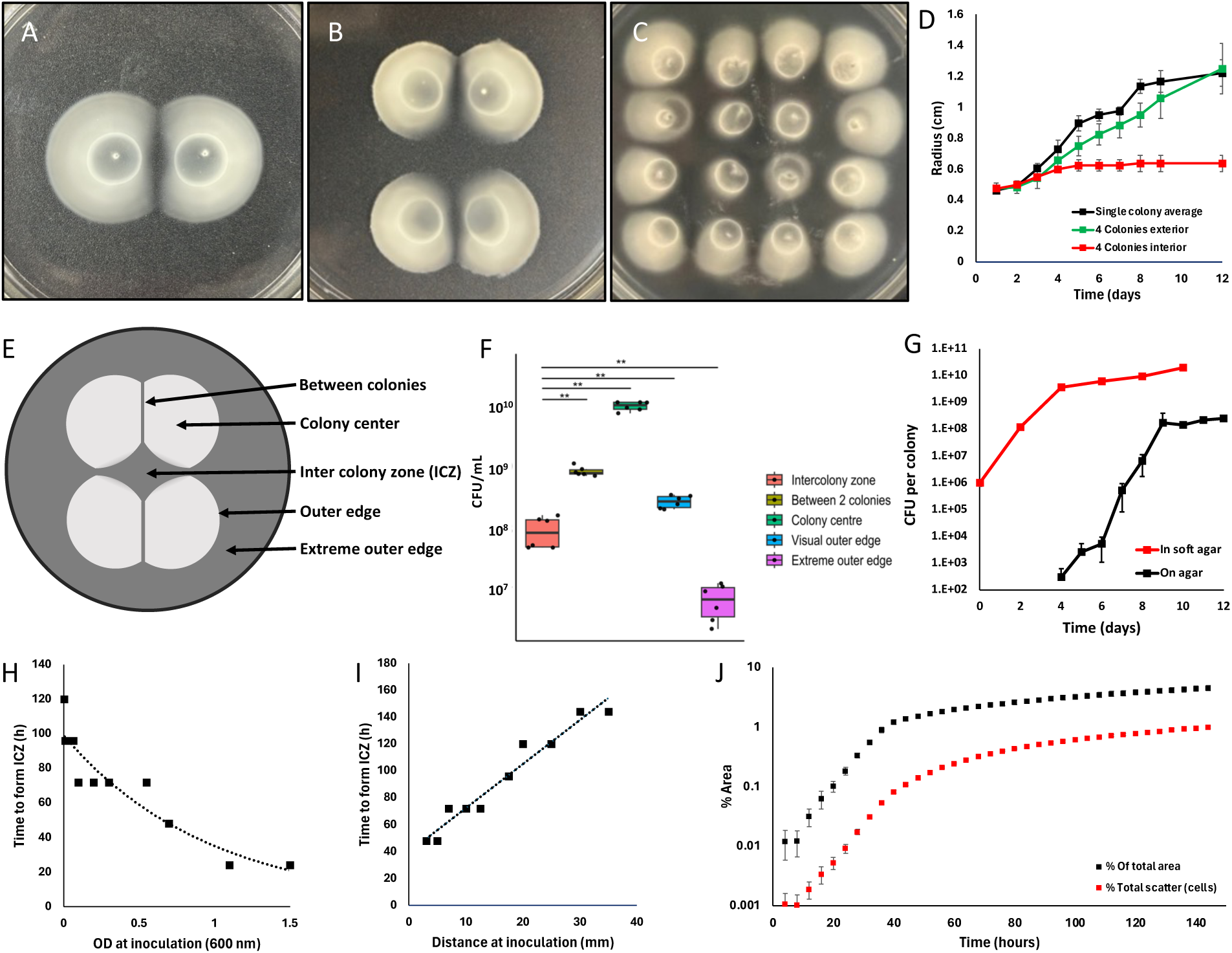
Sibling rivalry and colony arrest in *Bradyrhizobium diazoefficiens* USDA 110 are indicated by the formation of an intercolony zone (ICZ) between 2, 4, and 16 colonies on swim agar, demonstrating spatial patterns of colony arrest (A-C). The experimental setup used to investigate ICZ and colony interactions is shown as a schematic diagram (E). The number of culturable cells per zone, highlighting differences in cell viability and density within the ICZ compared to other areas (F); the culturable count of single colonies growing on 1.5% agar vs. the total biomass of a four-colony setup (G); colony expansion over time, illustrating dynamic changes in colony size and ICZ formation (D); the time needed to form the ICZ as a function of initial cell density (H); and the distance between colonies, providing insights into ICZ kinetics (I). Panel J shows the surface area of all colonies and the cell density as a function of light scatter in a four-colony setup (J).

### Cell density quantification across colonies

To determine the density of culturable cells across the colony, we partitioned the colony into five specific locations: 1) Inter-colony zone (ICZ), 2) the area between two colonies, 3) the colony center, 4) the outer edge, and 5) the extreme outer edge (Fig. 1E). We collected 100 µl of agar from each of the five locations into separate 1.5 ml tubes. To each tube, 900 µl of sterile water was added, and the suspensions were thoroughly mixed by vortexing for 30 s, shaking for 10 min, and vortexing for an additional 30 s. Colony forming unit/ ml (CFU/ml) was determined by plating 20 µl droplets of ten-fold dilutions in triplicate onto PSY agar and incubating for 5 d.

### Expansion visualization

To quantify colony expansion and time taken to develop the demarcation line, 4-colony setup plates were incubated for 12 d, and the radius of expanding inner and outer edge was measured every 24 h. To visualize colony expansion, we used a scanner (Epson Perfection V370 Photo) to capture images every 4 h, starting from 0 h up to 140 h (∼6 d) of incubation. Images were compiled using the montage feature of ImageJ software [37]. Scanner images (Fig. S2) were opened in Fiji and converted from RGB to 32-bit grayscale [38]. The mean background was measured, and image stacks were made for each colony and an intensity threshold was applied at the minimum detectable amount of cells corresponding to the ring at h 4. The area occupied by the colonies was quantified by the number of pixels occupying the threshold and the number of normalized number of organisms was quantified by the intensity originating from light scatter within the threshold. Colonies started using absorbance levels from 0.005 to 1.5 were observed every 12 h to investigate the relationship between the time taken to develop the demarcation line and the concentration of the starter inoculum. To investigate the role of initial distance 20 µl at the absorbance of 0.100 nm was spotted from 3 mm to 35 mm apart and observed every 12 h.

### Growth inhibitory effect on other strains and species

To determine whether USDA 110 impacted the growth of other *Bradyrhizobium* strains and other bacterial species (Table S1), a 10 µl drop of the specific strain (absorbance of 0.100) was introduced onto the ICZ of a 10-day-old colony. Bacteria were also inoculated onto sterile PSY motility plates to determine colony expansion on their own.

### Space maintenance

To determine the distribution of cells within swimming colonies, we started colonies of bjGFP tagged USDA 110 onto motility agar in glass bottom MatTek dishes (35 mm petri dish, no. 1.5 cover glass, P35G-1.5-14-C). Widefield images of colonies at the ICZ boundary, colony center, and colony outer edge were captured using an inverted microscope built around a Till iMic (Till Photonics, Munich, Germany) equipped with a 60X 1.2 N.A. water-immersion objective lens. Imaging was done from 237 µm to 321 µm above the coverslip with a step size of 2 µm at excitation/emission of 488/510 nm and 560/618 nm and for bjGFP and mChe respectively. A Z stack was captured for each fluorophore. The resulting image sequence was transferred to FIJI, and images from three different depths (245, 271, and 299 um) were shortlisted [38]. Image background was subtracted using the rolling ball background subtraction method with a ball radius of 25 pixels. The noise was then reduced by processing the images with a despeckle median filter.

To confirm spatial separation, we cultivated mChe, bjGFP, sYFP2, and mTq2-tagged USDA 110 strains individually for 48 h. After dilution to absorbance ∼0.100, 1 ml of each was combined, and the co-culture was inoculated onto soft agar as a four-colony setup. After ten d of incubation, swimming colonies were viewed using the same microscope settings as described above with excitation/emission of 515/527 nm for sYFP2 and 434/474 nm for mTq2.

### Physiological conditions of the cells

To determine whether colony arrest was due to death of cells, we stained colonies by flooding plates with, BacLight (Molecular Probes; Kit L13152) [25], and viewed fluorescence after 15 min using a fluorescence stereomicroscope (Olympus SZX16). Images obtained using GFP and RFP filters were composited in ImageJ. To assess the respiratory activity of the cells across the colony, we conducted the TTC (2,3,5-triphenyltetrazolium chloride) assay [25]. In brief, a 0.01% TTC (MP Biomedicals, LLC, SR00767) solution was prepared in water, and 5 ml was poured over 10-day-old swimming colonies. After an incubation period of 5 h at 25° C, plates were observed and imaged using iPhone 14 pro. ATP content was measured using the BacTiter-Glo™ Microbial Cell Viability Assay Kit (Promega, Cat.G8230/2). Briefly, 500 µl of agar from each of the five locations was transferred in a 1.5 ml tube with 750 µl of P/10. P/ 10 is 1/10 the peptone and yeast extract in PSY, with no sugar added [39]. The suspension was mixed by pipetting, vortexing for 30 s, shaking for 10 min, and vortexing again for 30 s. Subsequently, 100 µl of the suspension was used to determine the CFU/ml count. After centrifugation at 30,000 *x g* for 20 min at 4° C, the pellet was suspended in P/10 to a total volume of 500 µl, 250 µl of ice-cold 1.2 M perchloric acid was added, and the tubes were vortexed for 10 s and then incubated on ice for 15 min. After centrifugation at 30,000 *x g* for 7 min at 4° C, 500 µl of the supernatant was added to 250 µl of a mixture of KOH (0.72 M) and KHCO_3_ (0.16 M) was added to neutralize it. Following centrifugation at 30,000 *x g* for 10 min at 4° C, 100 µl of the supernatant was transferred to a well of a 96-well glass plate, and 100 µl of Bac Titer-Glo reagent (room temperature) was added, then shaken for 5 min in the dark. The reaction mixture was transferred to a 96-well white plate, and luminescence was read using a luminometer (BioTek Synergy2 Microplate reader). Relative Luminescence Units (RLU) were normalized by CFU/ml obtained from each sample location.

We wanted to assess the movement ability of cells in the Inter-Colony Zone (ICZ), colony center, and the extreme outer edge of the swimming colony. Movement assay was performed as described previously [39]. Briefly, agar from each of these locations was pooled to 7 ml and resuspended in 3.5 ml of sterile P/10 medium in 50 ml tubes. The suspension was mixed by vortexing for 30 s, shaking for 10 min, and vortexing again for 30 s. Suspension (3 ml) was added into 3 wells of a 12-well plate. The remaining sample was used to determine the initial CFU. Three glass capillaries filled with PSY arabinose (chemoattractant) were introduced into each well, and the plate was left for one hour to allow the cells to move toward the attractant [39]. After 60 min, the capillaries were transferred to microcentrifuge tubes, and the liquid was expelled from the capillaries. The expelled liquid was used to determine CFU of motile cells. The proportion of cells moving towards the attractant was calculated using initial and motile cell CFU. To gain insights into the role of the two known flagellar systems of USDA 110, colony expansion in a 4-colony-setup of wild type was compared to the delta-*fliC* strain, which lacks subpolar flagella, the delta-*lafA* strain, which lacks lateral flagella, and the delta-*fliC*/delta-*lafA* strain [40].

To investigate the repellent ability of the ICZ region, we pooled 7 ml of agar from the ICZ, exposed it to UV for 30 min to kill cells, and resuspended it in 3.5 ml of P/10 or PSY liquid medium. The tubes were shaken as described above, and the resulting liquid was transferred to three capillaries, representing a single technical replicate of a biological replicate. Chemotaxis toward the capillary content was determined by inserting capillary ends in a 3.5 ml culture of USDA 110 bjGFP-tagged cells in the well of a 12-well plate. The attraction of cells towards the ICZ with and without arabinose added was quantified by transferring the liquid from the capillaries to a 96-well plate and measuring fluorescence using a FLUOstar Omega reader (BMG LABTECH) at the excitation and emission maxima of 497/509 nm. To confirm the efficacy of UV treatment in eliminating all ICZ cells, capillaries containing the UV-treated ICZ samples suspended in attractant were dipped into wells that did not contain any culture.

### What could cause arrest in colony expansion?

To investigate whether nutrient depletion caused colony arrest, we supplemented the ICZ and the area between two colonies with 30 µl of ten-fold PSY medium on the 4th and again 7th d of the 10-d incubation period. Plates were observed every day, and images were recorded to monitor any changes in colony expansion. In parallel, we decreased the time for colonies to meet, decreasing agar density to 0.15%, as this allows faster colony migration and a shorter time to reduce total nutrients consumed. Plates were observed every day. To visualize the role of compression waves as a possible factor for ICZ development [27], we placed a sterile glass slide vertically between two swimming colonies and observed daily for any demarcation line development between the swimming colony and the glass slide. In a separate experiment, we inoculated colonies closer to the edge of the Petri plate, and colonies were expanded near the plate’s edge daily for demarcation line development. The rationale was that if generated, compression waves from each colony would interact with the glass slide or petri plate edge and arrest the movement of the cells at the colony edge.

To investigate whether colony arrest was due to a diffusible compound, we used a permeable membrane that allowed diffusion but not cell passage. We placed an autoclaved 0.2µm pore size filter membrane (Isopore PC Membrane, LOT:0000283982) over a 10-d-old bjGFP expressing a single swimming colony and a set of two swimming colonies (Fig. 5A). ICZ extracted from 5 plates was placed onto fresh motility plates, overlayed with membrane and soft agar, and spot inoculated (Fig. 5F). This procedure was repeated using autoclaved and UV-treated ICZ. We poured 2 ml of soft agar onto the center of the filter membrane, and after solidifying, we spot-inoculated with mChe expressing liquid inoculum (10 µl) at an absorbance of ∼0.100. The plates were observed for growth and motility of second drop inoculation (mChe) and imaged after 7 d of incubation. To evaluate the effect of lower colony on growth above in absence of potential dilution, we used membrane that allows only water to pass (Sterlitech, Dow Filmtec Flat Sheet Membrane, Batch: 67758). As an additional confirmation for the presence of a diffusible compound, transwells with membrane with a pore size of 0.4 µm (Corning Incorporated costar3450-clear) was placed over a 10-day-old 4-colony setup and a sterile plate. Fresh liquid culture (300 µl, absorbance set at 0.035 at 600 nm) was added to the transwell. The plates were wrapped with Parafilm and incubated in a moist container for 48 h when CFU/ml was determined. To investigate the potential involvement of quorum sensing and its associated molecule Isovaleryl-homoserine lactone (IV HSL) as effector, we acquired the *bjaI* mutant (AL17) from Dr. Caroline Harwood [41]. This strain was inoculated as a 4-colony setup alongside the parental wild-type strain USDA 110 spc4, and colony expansion was observed over a period of 10 d.

**Figure 2.**
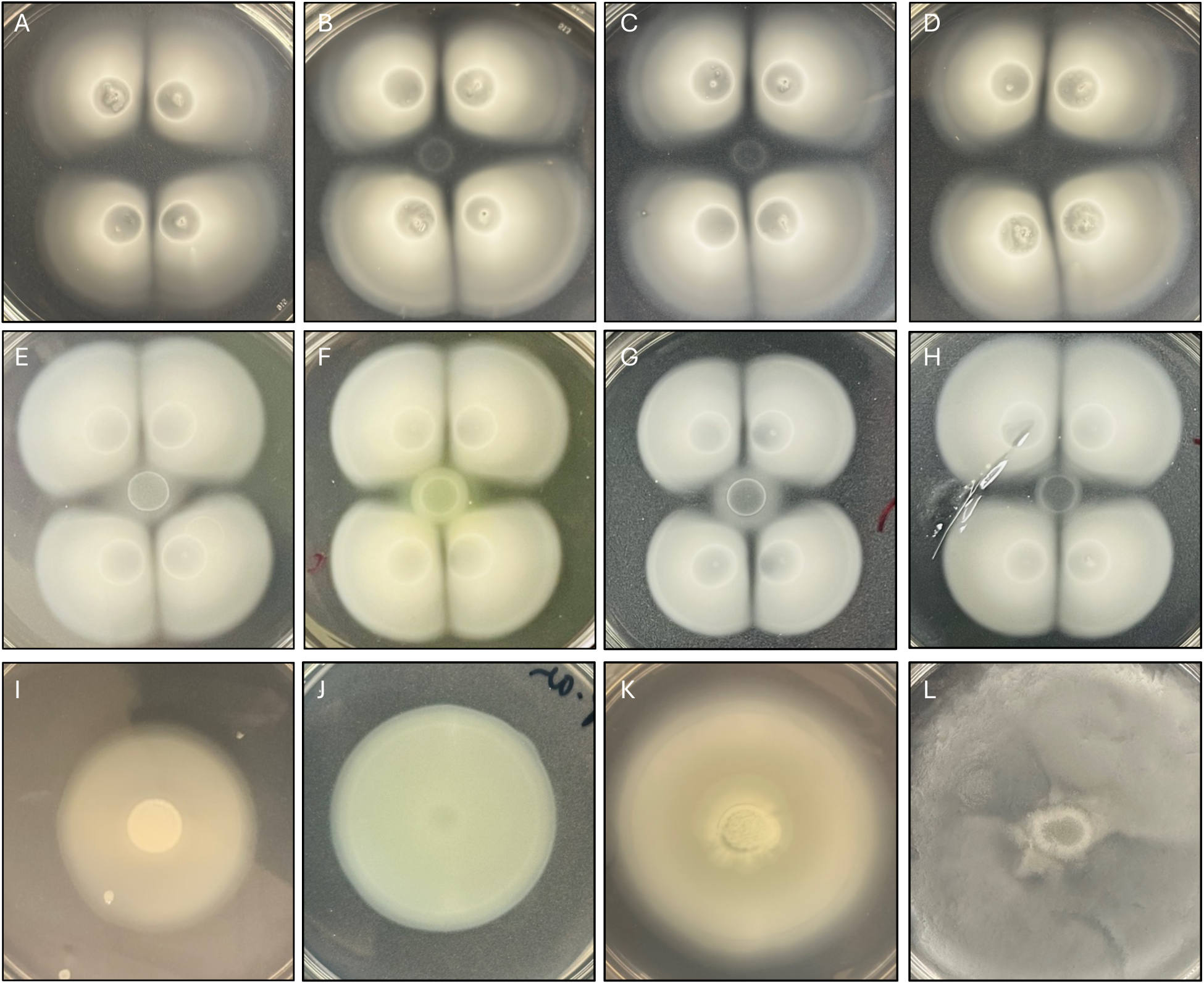
Inhibition of growth in the ICZ of swimming colonies of *B. diazoefficiens* USDA 110. Panels A-D were spot-inoculated onto the ICZ with USDA 110, USDA 20, USDA 140, and USDA 83. Panels E - H were spot-inoculated onto the ICZ with *Streptomyces ATCC*, *P. aeruginosa PA01*, *Herbaspirillum seropedicaea*, and *B. subtilis 168*, while panels I - L contained only the respective single cultures.

**Figure 3.**
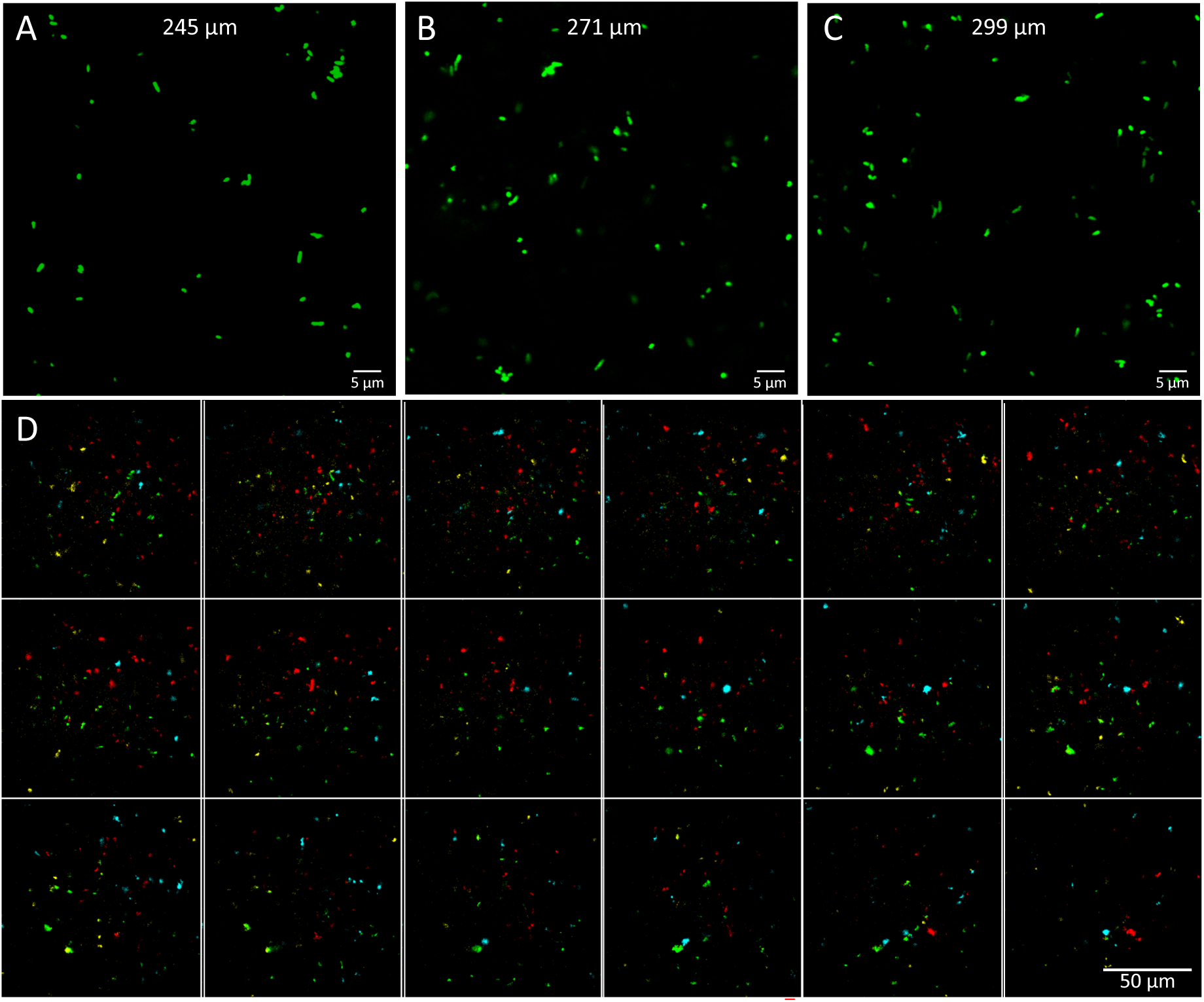
Distribution of USDA110 in soft agar viewed by confocal microscopy. Panels A - C show the cell distribution of bjGFP-labeled USDA 110 at 245, 271 and 299 µm. Panel D shows the distribution of four USDA 110 clones tagged with bjGFP, mChe, sYFP2 and mTq2, precultured separately and mixed in equal proportions before inoculation.

**Figure 4.**
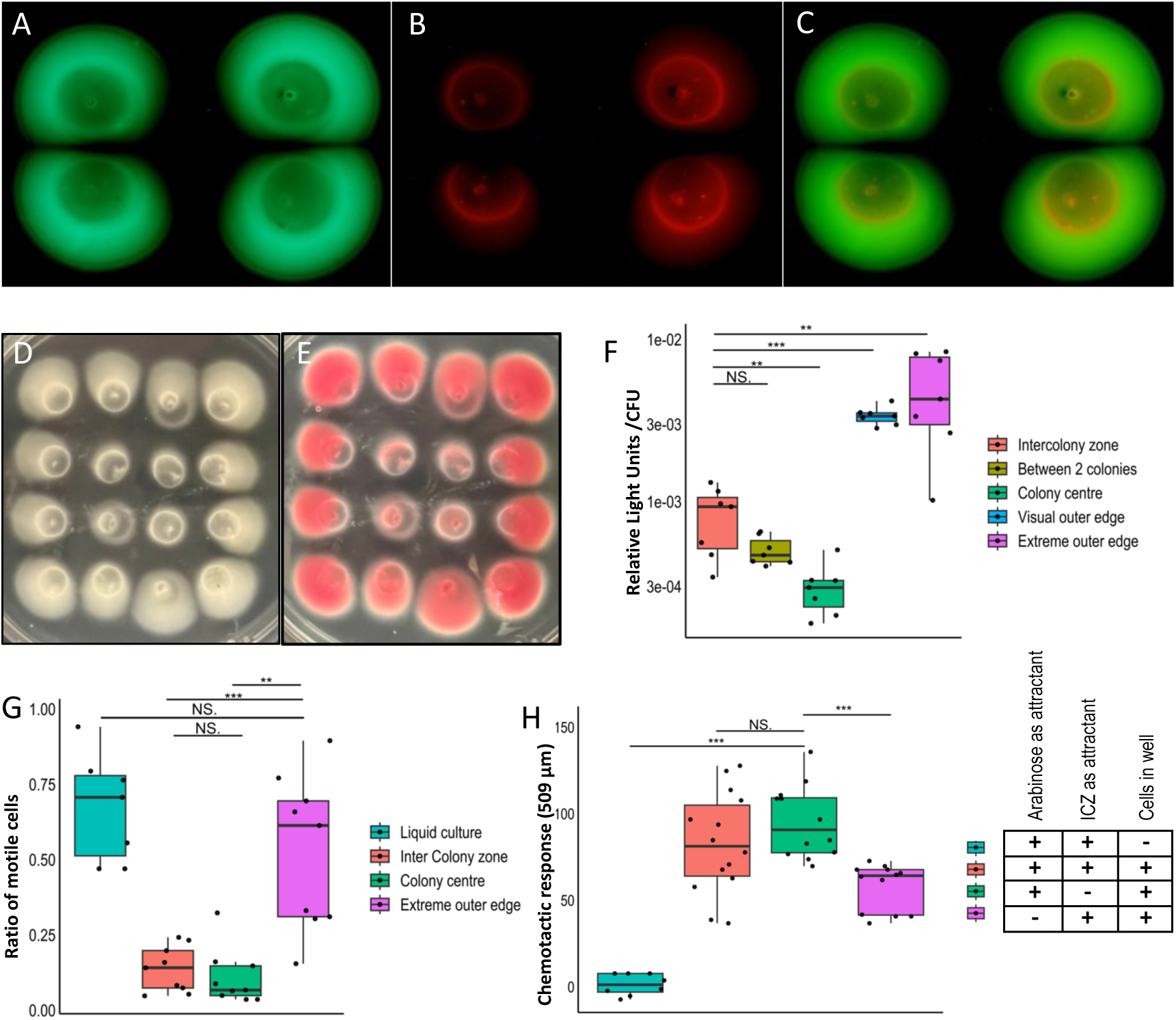
Physiological state of cells across swimming colonies of USDA 110. Panels A - C show four colonies stained with SYTO 9 and propidium iodide and merged to differentiate live (green) and dead (red) cells. Panels D - E show a sixteen-colony setup before and after staining with TTC, which turns pink through reduction linked to electron transport, indicating respiratory activity. Panel F shows ATP per cell in the five locations. Panel G shows the motility of cells taken from different locations toward arabinose compared to liquid culture. Panel H shows the chemotactic response of USDA 110 to ICZ, arabinose or both, as quantified by fluorescence.

**Figure 5.**
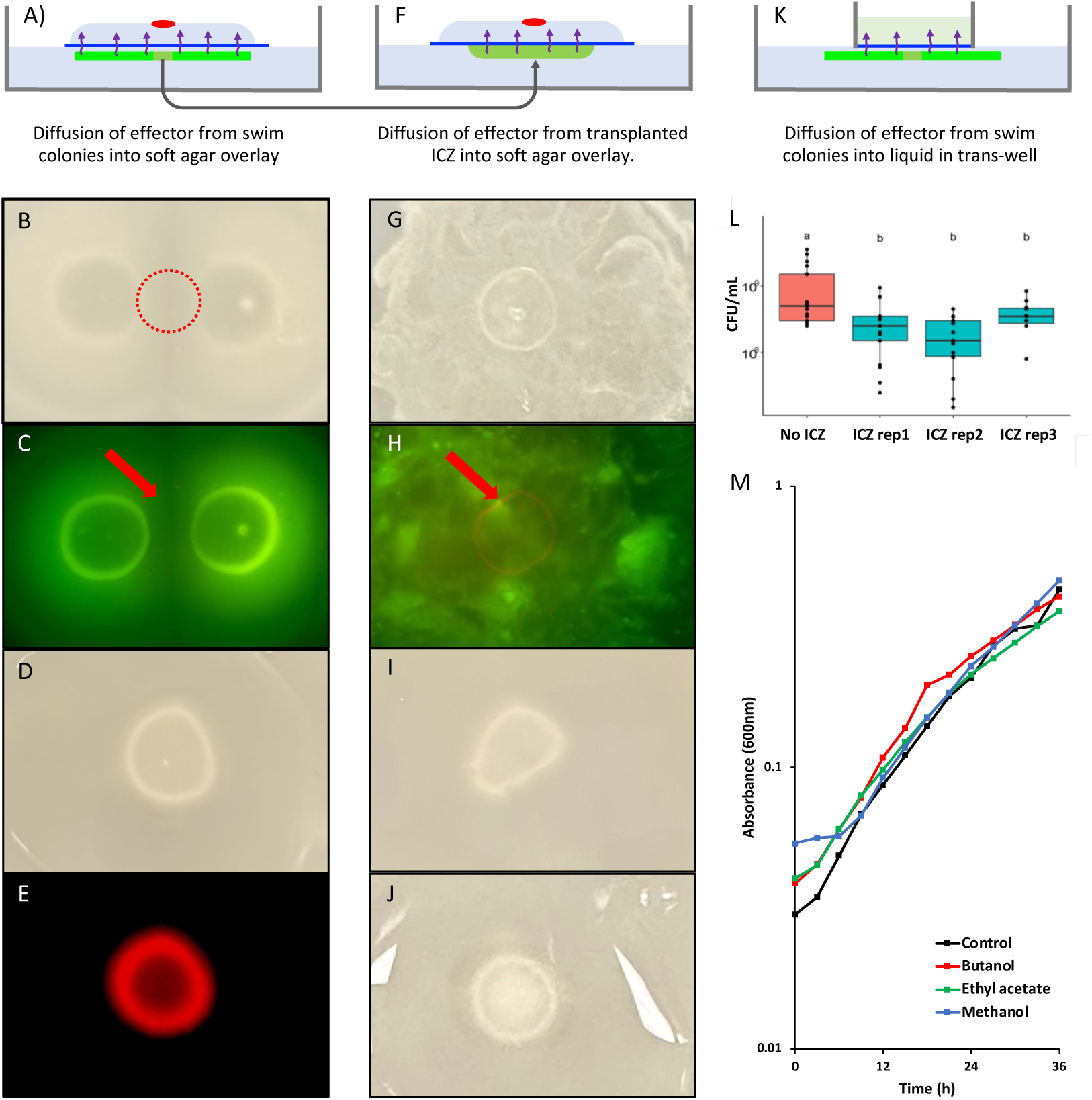
Experiments aimed at characterizing the cause of growth inhibition. Panels A, F and K illustrate the three experimental setups, with a blue line representing a membrane allowing the passage of molecules. Panels B–C show mChe-tagged USDA 110 introduced to soft agar over a 10-d-old 2 colony setup as outlined in panel a. Panels D–E show mChe USDA 110 culture introduced to soft agar over a plate with no underlying culture. Panels G and H show mChe USDA 110 culture introduced to soft agar over a layer of ICZ and transferred onto sterile soft agar, as outlined in panel F. Panel I shows USDA 110 introduced to soft agar over a layer of UV-treated ICZ, while panel J shows overlay onto autoclaved ICZ. Panel L shows the culturable count of USDA 110 in liquid medium in a transwell plate (0.4 µm pore size) incubated on four 10-day-old colony plates vs noninoculated plates. Panel M shows growth in liquid supplemented with extracts from ICZ using different solvents.

To extract the potential effector from the ICZ region, the agar between the 4 colonies “ICZ region” was collected from 40 plates and pooled into 50 ml tubes yielding 15 ml. Four such tubes were prepared to perform the extraction. The 15 ml extracted ICZ was suspended in 30 ml of either ethyl acetate, methanol, or water. The tubes were vortexed for 1 min and ultrasonicated with 5 pulses of 1 min each with intermittent cooling on ice for 30 sec between each pulse to avoid overheating of the sample. After sonication, the tubes were centrifuged at 7000 *x g* for 15 min, and the resulting supernatant, which was expected to contain the extracted chemical, was aliquoted into 2 ml tubes and concentrated in a speed vac for 3-4 h. Once dry, the tubes were refilled with 2 ml extract twice to increase the quantity of concentrate. The concentrated extract was stored at 4 °C until used. The extracts were individually resuspended in 100 µl of water, but the extract obtained from ethyl acetate was suspended in a 10% ethyl acetate solution in water. These suspensions were applied as 5 µl droplets at the edges of a 4-d-old young swimming colony, and the plates were observed daily for an additional 6 d period to observe a halt in colony expansion. To examine the impact of the extracts on the growth rate of USDA 110, 20 µl of the extracts were introduced into 150 µl fresh culture, and the cultures were inoculated as replicates of 9 in 96-well plates. Growth was monitored for 36 h in a plate reader (FLUOstar Omega reader (BMG LABTECH)) at 600 nm at 28 °C. To detect any protein occurring in ICZ, 30 ml of pooled ICZ was suspended with 15 ml of sterile water by vortexing and shaking steps [25]. Briefly, samples were frozen at −80°C for 2 h, thawed, and centrifuged (30000 *x g* at 4° C for 20 min) to remove pellet-containing cells and debris. The supernatant was subjected to TCA and ammonium sulfate precipitation at 80 % saturation, and the precipitate was harvested by centrifugation and suspended in PBS. The resulting precipitate was placed in a spin column with 3 kDa cut off (thermos scientific, Pierce Concentrator, PES, lot: XA337655) and centrifuged at 30000 *x g* for 10 min. The concentrate was prepared for electrophoresis and resolved in a 12% SDS PAGE gel. Gels were stained with Coomassie Brilliant Blue R250, destained, and viewed. We also employed SYPRO Orange Protein stain (Millipore Sigma), a protein dye, on the ICZ to look for any fluorescence under a stereomicroscope.

### Statistical Analysis

Experiments were conducted in triplicate biological replicates, each with three technical replicates. Fisher’s LSD test were performed in R to analyze data to determine statistically significant differences between the means of different groups [42]. Asterisks are used to denote levels of significance in p-values as: p < 0.05; *p < 0.01: **; p < 0.001: ***. R packages used included dplyr, agricolae, ggplot2, ggpubr, scales, readxl and patchwork.

## Results

### Characterization of the biological phenomenon

The development of a demarcation line between encroaching swimming or swarming colonies supporting is indicative of spatial inhibition and has been observed for different strains of a species [43]. However, sibling rivalry (competition between daughter colonies) is rarely documented and studied. Here we observed that identical strains of *Bradyrhizobium* develop a demarcation line when swimming colonies are inoculated next to each other, indicating that each colony prevents the expansion of the other (Fig. 1A). This behavior was observed in all five motile strains evaluated: *B. diazoefficiens* USDA 110*, B. elkanii* USDA 83*, B. japonicum* USDA 20*, B. arachidis* USDA 3384, and *B. liaoningense* USDA 13 (Fig. S1). We chose to characterize this phenomenon using *B. diazoefficiens* USDA 110, an agriculturally significant strain that fixes nitrogen in nodules of soybean roots. The observed behavior is not specific to the growth medium, occurring both in PSY-arabinose and SESOM soft agar. Four colonies had a more pronounced inhibition zone, which we termed “inter-colony-zone” (ICZ) (Fig. 1B). The colonies at the center of a 4 x 4 grid were suppressed the most, and the outer-most colonies showed expansion at their exterior edge and colony arrest at inner edges facing other colonies (Fig. 1C). The inhibition zone became more distinct with an increase in the number of inoculation points, indicating that the expansion arrest is related to regional cell density. For most of the experiments in this study, we standardized the inoculation setup to 4 colonies (Fig 1E).

To see if cells were evenly distributed across colonies, we determined the culturable count at five different locations of the 4-colony system (Fig. 1F). As anticipated, the colony center contained the highest number of cells, followed by the area between the two colonies (demarcation line) and the visual outer edge. Although the agar in regions outside of colonies appeared clear, ∼10^7^ CFU/ml were detected, 4 mm away from the visible colony boundary. This suggests that a subset of the colony population swam ahead, behaving like scout cells (Fig. 1F). Similarly, the occurrence of ∼10^8^ CFU/ml of cells in the ICZ suggested that a subset of the colony population swimming inward was suppressed and did not attain full density after one month of incubation. The presence of cells in the extreme outer edge and the ICZ was confirmed by fluorescence microscopy of GFP-tagged cells.

To characterize colony expansion arrest, we tracked in- and outward facing radii of the 4-colony setup vs single colony. Colonies expanded in all directions till day 4, when the arrest of movement towards each other became apparent because the inward-facing radius no longer increased (Fig. 1D and S2). The outward colony expansion at the 4-colony exteriors followed the same rate as that of a single colony.

To determine whether colony arrest is unique to swimming colonies, we compared the growth rate of cells in 4-colony setups (0.35%) versus colonies developing from single cells on solid agar (1.5%) through CFU counting. Colonies developing from single cells became visible after 4 d, hence these data start from d 4 (Fig. 1G). The generation time of cells in swim colonies was 8.1 h during the first 4 d, slowing down to 59.5 h from d 4 onwards. Cells in colonies on hard agar grew faster, dividing every 6.3 h till d nine when generation time slowed to 123 h, essentially stationary phase (Fig. 1G). The generation time of USDA 110 was inherently slower in soft agar, indicating some growth inhibition from the start. We also confirmed growth suppression by quantifying cell biomass of scanned colonies over time. After 40 h the rate of colony expansion decreased (Fig. 1J). Importantly, the rate of increase of biomass also decreased after 40 h, as reflected by the pixel number per colony. The discrepancy between onset of slower growth rate (Fig. 1G) and biomass increase (fig. 1J) may be because of the cells becoming smaller between 40 and 96h.

The initial cell number inoculated for founder colonies affected the time taken to ICZ (Fig. 1H). This indicated that demarcation lines form irrespective of initial inoculum density, but more time is required with a lower initial inoculum to reach the cell density required for colony arrest. Distance between founder colonies appeared linearly related to time for ICZ development (Fig. 1I). The greater the initial distance, the more time was required for expanding colonies to reach proximity. Collectively, these data show that ICZ formation is a function of cell density and proximity of colonies.

### Growth inhibitory effect on other strains and species

To determine whether reduced cell density in the ICZ was due to growth suppression, we introduced a fresh inoculum of USDA 110 to the ICZ of a 10-d-old 4-colony setup. Even after 10 d of incubation, no visible growth was observed (Fig. 2 A). Suggesting the presence indeed of a growth-suppressing phenomenon associated with swimming colonies of USDA 110. To determine whether other *Bradyrhizobium* strains and soil bacterial species were suppressed, we added cell suspensions of USDA 26, USDA 140, and USDA 83, and *B. subtilis*, *Salmonella* Typhimurium, *Herbaspirilum seropedicae* ATCC 33892, *Streptomyces* ATCC 49182, *Arthrobacter aurescens* TCI, *P. aeruginosa* PA0, *Pseudomonas* ADP and *E. coli* K 12 to the ICZ. All *Bradyrhizobium* strains were strongly inhibited, but USDA 26, USDA 140, and USDA 83 exhibited some growth within the ICZ (Fig. 2 B-D). Growth and motility of all the eight soil species were suppressed when compared to swimming on their own, but only 4 of these are shown (Fig. 2 E-L). The strongest impact on colony expansion was observed with *B. subtilis,* but even the aggressive *P. aeruginosa* was suppressed. This indicates that the growth-suppressing phenomenon affects not only USDA 110 but also other strains of *Bradyrhizobium* and other fast-growing species of soil bacteria.

### Space maintenance among cells of Bradyrhizobium

To determine the cell distribution of bjGFP tagged USDA 110 in the swimming colonies in soft agar, the ICZ, and the outer edge we performed confocal microscopy at 600x. Cells appeared distributed throughout the depth of the colony either as single or small groups, with no larger groups or clumps viewed at three depths of the motility agar (Fig. 3A-C). Similar observations were made for cells in the ICZ and the outer edge. While bacteria are widely known to form multicellular aggregates, both on surfaces and in swim agar [44; 45], USDA 110 displayed spatial separation among cells. This points to an ability to maintain spatial separation. To verify active spatial separation, we prepared separate exponential phase cultures of USDA 110 tagged with either mChe, bjGFP, sYFP2 or mTq2, mixed at equal ratios and co-inoculated onto soft agar. After 10d incubation, confocal microscopy revealed distinct separation among cells and small groups of the 4 colors throughout the agar (Fig. 3D). This separation between isogenic populations indicated the maintenance of spatial segregation and suggests a remarkable capability to distinguish its isogenic self when precultured populations are brought together. Distinct entities remained separate even when forced to grow in the same environment for 10 d. Notably, this segregation was also observed in non-swimming colonies of a mixed culture of mChe and bjGFP on hard agar, where they do not even have any space to move or expand away from one another (Fig. S3). These results show that while most cultured bacteria tend to coalesce, *Bradyrhizobium* maintains a distance among cells.

### Physiological state of the cells in the ICZ

We sought to determine the physiological state of the cells across colonies and in the ICZ. BacLight was used to visualize live versus dead cells, TTC reduction was used to visualize respiratory activity, and cellular ATP content was quantified. Baclight stained colonies fluoresced green throughout (Fig. 4 A). Only the circumference of the initial area of inoculation fluoresced red, indicating dead cells at the origin of the colony (Fig. 4B and C). No red fluorescence was observed at the edge of the colony, even the part facing the ICZ. This indicated that the observed halting of colony expansion was not due to loss of cell viability. This was further supported by the culturability of cells from the ICZ (Fig. 1F). Furthermore, the transfer of sections of the ICZ onto sterile agar results in visible growth after incubation. Populations closer to the ICZ and colony edge displayed less TTC reduction than at the colony center (Fig. 4D and E). Colorless TTC 2,3,5-triphenyl tetrazolium chloride is reduced to pink 1,3,5-triphenylformazan (TPF) by the activity of succinate dehydrogenase, a function of respiratory activity [46]. This indicated that cells in between colonies and at the colony’s outer edge showed lower respiration activity than at the colony center. The greatest suppression of respiration was observed in the s

4 colonies situated in the center of the 4×4 grid, characterized by minimal colony expansion and a light pink color with areas exhibiting a pronounced white tint. The amount of ATP per cell was significantly different across the colonies (Fig. 4F). Intriguingly, cells in the colony center had the least ATP, followed by cells in between the two colonies and the ICZ. Cells at the outer and extreme outer edge had the most intracellular ATP. Cells at the colony center displayed high respiratory activity but low ATP, indicating high energy demands. Cells between two colonies had lower respiration and ATP, indicating lower activity but high energy demands. Conversely, cells at the outer edge had lower respiration but high ATP, suggesting lower demands on cells.

To determine the motility status of the cells across the colony, we quantified the motility of cells towards the attractant arabinose. While populations at the outer edge displayed motility similar to that of cells grown in liquid culture, populations in the ICZ and the colony center displayed significantly lower swimming (Fig. 4 G). This indicated that the observed phenomenon of colony arrest is also associated with a lack of motility. Because USDA 110 expresses at least two flagellar systems [47], we evaluated colony expansion by mutants deleted for either *lafA,* or *fliC*, or both. Both of the single knockouts showed colony expansion and colony arrest as the wild type, indicating that either of the flagellar systems support colony expansion on their own (Fig. S4). As sibling rivalry was observed in both single flagellar mutants, the inhibitory function is not linked to either system. The mutant lacking both flagellar systems did not swim. To determine if the ICZ acts as a chemorepellent, we conducted an attraction comparison between the agar from the ICZ and the known attractant, L-(+)-arabinose. There was no significant difference between swimming towards the ICZ and the attractant, indicating that the ICZ did not exhibit chemo-repellent properties (Fig. 4H). These results indicate that cells exposed to the ICZ lose motility and respiratory activity but are not chemotactically repelled but are not dead.

### Possible causative agent of colony arrest and growth suppression

To explore possible causes of expansion arrest and growth suppression, we explored hypotheses related to the depletion of nutrients, compression waves, and diffusible compounds or effectors. Assuming that decreased incubation time would lead to decreased nutrient consumption, we decreased agar density to 0.15% to decrease time required for expansion. Inoculation into 0.15% agar resulted in an accelerated expansion of colonies, and colony arrest and ICZ formation was observed after two rather than 4 d. Addition of droplets of 10 times higher concentration of nutrient to the ICZ four and seven d after inoculation had no noticeable impact on colony arrest (Fig. S5). We concluded that the cessation of movement in approaching colonies was not caused by nutrient depletion. As Espeso et al. proposed that the arrest of swimming populations of *P. putida* was due to the generation of compression waves that propagate through the medium towards the other colony, we examined compression waves as a potential factor [27]. We introduced a glass slide as a physical barrier between two expanding colonies, and inoculated colonies close to the edge of the petri plate. Compression waves would reach these barriers, resulting in the formation of a demarcation line. We observed that the edges of the colonies progressed to the glass and plastic boundaries with no visible demarcation line (Fig. S6). We concluded that the production of compression waves is not a contributing factor to the phenomenon of colony arrest in *Bradyrhizobium*.

To ascertain the possibility of a diffusible effector responsible for colony arrest, we employed 0.2 μm membrane filters to allow passage of diffusible compounds but not cells. Membrane filters were placed onto 10 d old two or single colony setups of bjGFP tagged cells, overlayed with fresh motility agar, and spot inoculated with mChe tagged cells to observe suppression or growth (Fig. 5A). Notably, the fresh inoculum on the top layer of agar did not develop into a visible colony, and no red fluorescence was detected (Fig. 5B and C). Similar observations were made with a single colony under the membrane. Growth above a membrane allowing only water to pass was uninhibited, suggesting that growth inhibition requires a diffusible compound and is not due to any compression wave action. The inoculum placed on top of a system, with no swimming colonies at the lower layer of agar (sterile), grew into visible colonies and fluoresced red (Fig. 5D and E). The absence of growth in the upper layer indicated growth suppression, supporting the presence of a growth-suppressing diffusible compound produced by the underlying swimming colonies. To evaluate the presence of diffusible compounds specifically in the ICZ, we removed ICZ from 10-d-old 4-colony setups and transferred it either untreated, UV treated, or autoclaved onto fresh motility agar, followed by membrane overlays above (Fig. 5F). Newly inoculated cultures grew only minimally, indicating the presence of diffusible compounds in the ICZ specifically (Fig. 5G and H). During incubation of untreated ICZ, visible growth appeared as seen by green fluorescence. We ascribed this to the low number of cells present in the ICZ (Fig. 1F) that started growing in the absence of their surrounding colonies. To exclude contributions from this new cell activity, cells in ICZ were killed by either UV or heat treatment. The new inoculum exhibited similar growth on both, as compared to the control without underlying ICZ (Fig. 5I and J). The lack of strong growth suppressing the impact of ICZ indicates that the effector has a short half-life or is sensitive to UV radiation and high temperature. The observation that growth suppression occurs over established colonies but far less over freshly outgrowing culture from ICZ indicates that the functioning of the effector is most active in the presence of live cells but has the ability to diffuse through agar and work at a distance. To assess growth inhibition by the effector of cells in a liquid culture, we placed a transwell with a pore size of 0.4 μm on 10-d-old colonies (Fig. 5K). Significantly lower yields were observed in transwells over swimming colonies than over sterile agar in three separate experiments, indicating growth suppression (Fig. 5L). This observation further reinforces the conclusion that a diffusible “Effector” is produced by swimming colonies of USDA 110, responsible for the growth suppression of new cultures.

With evidence for a diffusible effector, we sought to identify the nature of the compound responsible. We did not find a change in pH in the ICZ, indicating the absence of a strong organic acid or base as an effector. To evaluate the possible role of quorum sensing, we compared the colony expansion of the wild type versus a *bjaI* deletion mutant shown to be deficient in quorum sensing [41]. QS mutants behaved as the wild type, indicating that the known QS system of USDA 110 is not responsible for colony arrest. We attempted to extract a possible protein from the ICZ by TCA or ammonium sulfate precipitation and separately by concentrating using 3kDa cut-off spin filters. However, we were unable to detect any protein bands on SDS-PAGE gels. No fluorescence was detected after adding the protein-detecting SYPRO Orange Protein stain (Millipore Sigma). These results show that the effector is unlikely to be a protein larger than 10 kD, but if it is, it has a very short half-life. Unable to directly determine the nature of the effector, we performed extractions from ICZ using four solvents: water, butanol, ethyl acetate, and methanol. Application of concentrated extract (5 µl) at various distances surrounding young colonies did not lead to visible arrest of expansion (data not shown). Likewise, the growth of liquid cultures supplemented with concentrated extracts grew at the same rate as that of untreated cultures (Fig. 5M). These observations suggest that the effector has a low half-life or is lost in the extraction or concentration process.

## Discussion

We report self-growth suppression in *B. diazoefficiens* USDA 110, initially observed as suppression of colony expansion, but exhibiting suppression of cellular activity and motility by surrounding cells through sibling rivalry. While sibling rivalry has been previously reported in *M. algicola, P. putida,* and *P. dendritiformis* [25; 26; 27], *Bradyrhizobium* is unique as it suppresses growth of itself (sibling rivalry), related strains (kin rivalry), and a broad cross-section of other bacterial taxa (antagonism). We characterized this phenomenon and reported experiments to understand the underlying mechanism. Sibling rivalry is not unique to USDA 110, as it occurred across various *Bradyrhizobium* strains. The phenomenon also occurred in SESOM, an extract from soybean field soil, indicating occurrence in natural soil environments, thereby emphasizing the ecological relevance of expansion suppression beyond laboratory nutrition conditions.

Expansion suppression of *Bradyrhizobium* appears due to inhibition of cells rather than cell death. The generation time of cultures introduced into soft agar was significantly slower than on hard agar (8.1 *vs* 6.3h), indicating the onset of growth suppression from the beginning. The increase in population was associated with further slowing of average generation time in soft agar (∼59 h from 4^th^ d vs 6.3h till d 9 on hard agar). Cell movement and growth cease in *Myxococcus xanthus* at low nutrient levels [43]. The shift to the quasi-stationary phase in USDA 110 occurred despite the availability of nutrients and space between cells, indicating a different basis for inter-colony rivalry. The onset of stasis was associated with an increase in population and time of populations occurring in agar. Cells in the colonies had reduced ATP/cell as compared to those at the colony outer edge. Furthermore, colonies surrounded by others displayed a reduced respiration rate (TTC reduction). Thus, cells surrounded by other colonies display lower overall activity and fitness than cells facing unpopulated space. Likewise, cells in colonies and the ICZ displayed low motility as compared to cells facing unpopulated space. The cells in freshly populated space displayed higher activity due to distance from established populations. A similar decrease in motility and respiration was observed during the study of sibling rivalry in *M. algicola* [25]. Bacteria, in general, spread outward for constant supply of a better nutritional environment [48]. MDR *Salmonella enterica* serovar Typhimurium has reduced motility, both swimming and swarming, in the presence of antibiotics [49]. While sibling rivalry in *P. dendritiformis* was caused by the death of cells [26], USDA 110 populations at the rivalry phase were culturable and stained live by BacLight. These observations point to suppression rather than killing as the cause of colony expansion suppression. However, the suppressive effect was strong enough to inhibit the growth of fresh inoculum on established ICZ (Fig. 2A, 5B, C and L).

A subset of cells was found outside of the visible colony boundary, both facing other colonies (ICZ) and unpopulated space. While cells in the ICZ were metabolically suppressed and did not display swimming, cells at the outer edge swam actively. These cells behaved like scouts as they occurred far from the colony. This indicates a subset of the population displaying phenotypic heterogeneity. We have previously observed phenotypic heterogeneity in *B. diazoefficiens* USDA 110 while studying its surface properties [35]. Intriguingly, they did not grow locally to high density, indicating some restraint on cell division despite their ATP status. This swimming away from the colony could not be ascribed to chemorepulsion (Fig. 4H). The physiology of this subpopulation should be further studied.

Growth suppression by *Bradyrhizobium* appeared to be due to a diffusible compound with a short half-life. This effector of growth suppression was able to diffuse through the aqueous agar matrix and through pores in polycarbonate and polyester membranes. It lost activity after exposure to UV and heat, and during extraction and concentration procedures, indicating lability or short half. Sibling rivalry in *P. dendritiformis* is caused by a two-protein system of the pre-toxin Slf and protease subtilisin [29]. *M. algicola* sibling rivalry was associated with GDPD, with glycerophosphoryl diester phosphodiesterase as the predicted effector [25]. The USDA 110 effector is unlikely to be proteinaceous as we were unable to obtain an extracellular protein from the ICZ. The effector is also not a strong acid or base, as no shift in pH was observed. It does, however, accumulate over time and as a function of population size since colonies surrounded by multiple others were most suppressed (Fig. 4D and E). This behavior seems to be similar to the concentration-dependent mechanism of antibiotics [50]. The effector is clearly broad range, as it suppresses the growth of producers, members of the same genus, and a broad cross-section of other bacteria (Fig. 2). The apparent short half-life poses a challenge for the chemical characterization of the effector as the activity of extracts must be verifiable. This has prevented us from pursuing chemical characterization to date.

Self-growth suppression or sibling rivalry leads to curtailing of the population, which should limit the success of the species in soil. *Bradyrhizobium* are slow growing and have to compete with diverse faster growing bacteria in soil and the rhizosphere [51]. These self-supressing traits are counterintuitive in terms of long-term persistence and are expected to cause the species to be unsuccessful and therefore rare. Intriguingly, *Bradyrhizobium* is the most predominant genus found in soils globally [32], indicating the success of the species in soil. This indicates that self-growth suppression, as described here, is part of the success strategy used by *Bradyrhizobium*. Copiotrophs such as *Bacillus* and *Pseudomonas* devour resources to maximize their population, outcompeting slower growing species [52; 53; 54]. Because nutrient input into soil is sporadic or seasonal, copiotrophs devour these limited nutrients, causing the community to starve [55; 56]. The ability of *Bradyrhizobium* to restrain or suppress growth and activity of itself and copiotrophs should reduce nutrient consumption, causing longer-term nutrient availability. This would enable the species to have longer-term success. We, therefore, hypothesize that the self-suppressing mechanism indicates a strategic adaptation for conserving energy and resources in a competitive and nutrient-limited soil environment. This concept is similar to entering a dormant state and using fewer resources, giving it an edge over copiotrophs, especially in oligotrophic soil environments [57]. A similar strategy has previously been reported for bacteria growing slowly or becoming dormant under starvation conditions in soil as an alternative survival strategy [58]. By strongly suppressing incoming bacteria, an individual *Bradyrhizobium* strain would dominate its location, facilitating its colonization of spaces such as root surfaces. This would benefit colonization, including nodulation of legume hosts [59]. We relate self-growth suppression to the long-term survival and competition strategy of this agriculturally beneficial genus.

## Acknowledgments

We thank Dr. Nicholas Butzin for their critical reading of the manuscript, and Dr. Daniel Petras for his guidance on extraction of putative compounds. We would also like to extend our thanks to Dr. Caroline Harwood, Dr. Hans-Martin Fischer, Dr. Brittany Belin, Dr. Aníbal Lodeira and Dr. Michael Sadowsky for providing us with strains, mutants and genetic constructs.

## Author Contributions

Volker Brözel and Armaan Sandhu conceptualized the project. Armaan Sandhu, Volker Brözel, Adam Hoppe and Brady Fischer analyzed the data. Senthil Subramanian and Volker Brözel acquired the funding. Armaan Sandhu, Volker Brözel, Adam Hoppe and Brady Fischer performed the experiments. Armaan Sandhu, Volker Brözel, Adam Hoppe and Brady Fischer developed methodology. Armaan Sandhu and Volker Brözel wrote the manuscript. Armaan Sandhu, Volker Brözel, Adam Hoppe, Brady Fischer and Senthil Subramanian reviewed and edited the manuscript.

## Supplementary material

Supplementary material is available at The ISME Journal online.

## Data availability

All the data generated for analysis during this study are included in this published article and its supplementary information files.

