## Supplementary material for "Self-growth suppression in *Bradyrhizobium diazoefficiens* is caused by a diffusible antagonist": Table S1

Table S1 Strains and plasmids used in this study

| Bacterial strain | Source | Reference |
|---|---|---|
| *Arthrobacter aurescens* TC1 | Dr. Mike Sadowsky | [1] |
| *Bradyrhizobium. arachidis* USDA 3384 | NRRL * | |
| *B. diazoefficiens* USDA 110 | NRRL | |
| *B. diazoefficiens* USDA 110 *spc4* | Dr. Caroline Harwood | [2] |
| *B. diazoefficiens* USDA 110 spc4 *fliC,* devoid of subpolar flagellar filaments | Dr. Aníbal R. Lodeiro | [3] |
| *B. diazoefficiens* USDA 110 spc4 *laf A* and *fliC,* lacking both flagellar systems | Dr. Aníbal R. Lodeiro | [3] |
| *B. diazoefficiens* USDA 110 spc4 *lafA,* devoid of lateral flagellar filaments | Dr. Aníbal R. Lodeiro | [3] |
| *B. elkanii* USDA 26 | NRRL | |
| *B. elkanii* USDA 83 | NRRL | |
| *B. japonicum***** USDA 110 spc4 *bjaI* mutant (AL17) | Dr. Caroline Harwood | [2] |
| *B. japonicum* USDA 126 | NRRL | |
| *B. japonicum* USDA 140 | NRRL | |
| *B. japonicum* USDA 20 | NRRL | |
| *B. japonicum* USDA 6 | NRRL | |
| *B. liaoningense* USDA 13 | NRRL | |
| *Bacillus subtilis* 168 | BGSC** | |
| *Escherichia coli* K12 | ATCC | |
| *E. coli* S17-1 λpir | Nova lifetech Inc | |
| *Herbaspirillum seropedicae* ATCC 33892 | ATCC*** | |
| *Pseudomonas ADP* | Dr. Mike Sadowsky | [4] |
| *P. aeruginosa* PA0 | ATCC | |
| *Salmonella* Typhimurium | Lab collection | |
| *Streptomyces* ATCC 49182 | ATCC | |
| Plasmids used | | |
| pRJPaph-bjGFP-1 | Dr. Hans-Martin Fischer | [5] |
| pRJPaph-mTq2-1 | Dr. Hans-Martin Fischer | [5] |
| pRJPaph-sYFP2-1 | Dr. Hans-Martin Fischer | [5] |
| pRJPaph-mChe-1 | Dr. Hans-Martin Fischer | [5] |

*NRRL Culture Collection of the Agricultural Research Service, USDA

** Bacillus Genetic stock Center
*** American Type Culture Collection
**** Till 2013 *B. diazoefficiens* USDA 110 was known as *B. japonicum* USDA 110[6]
