## Supplementary material for "Self-growth suppression in *Bradyrhizobium diazoefficiens* is caused by a diffusible antagonist": Figures S1 - 6

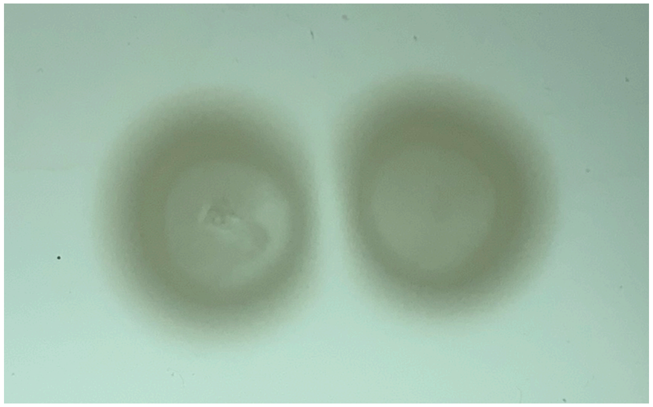

**USDA 13**

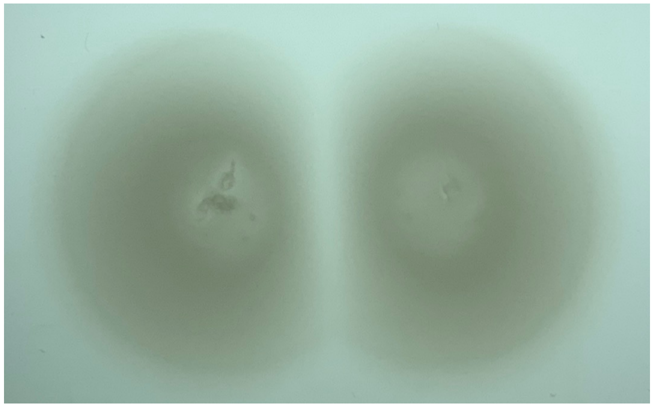

**USDA 20**

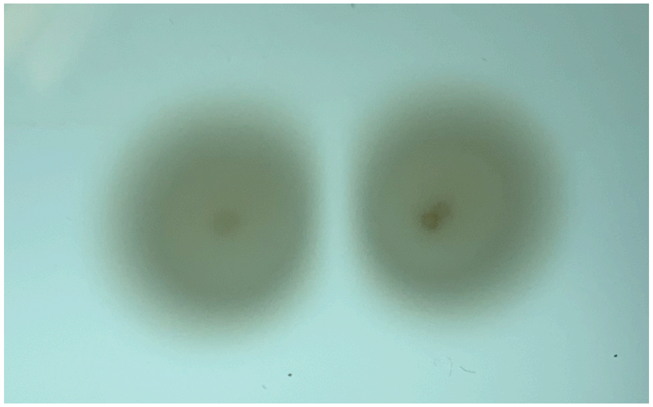

**USDA 83**

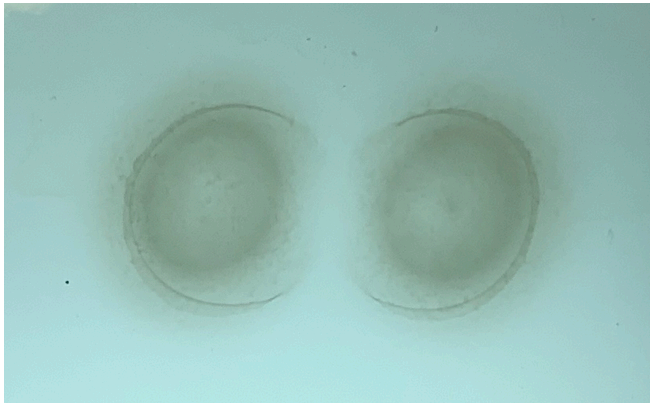

**USDA 3384**

Figure S1 Two colony set-up for *Bradyrhizobium* USDA 13, 20, 83 and 3384 in PSY soft agar with arabinose, incubated for 7 d.

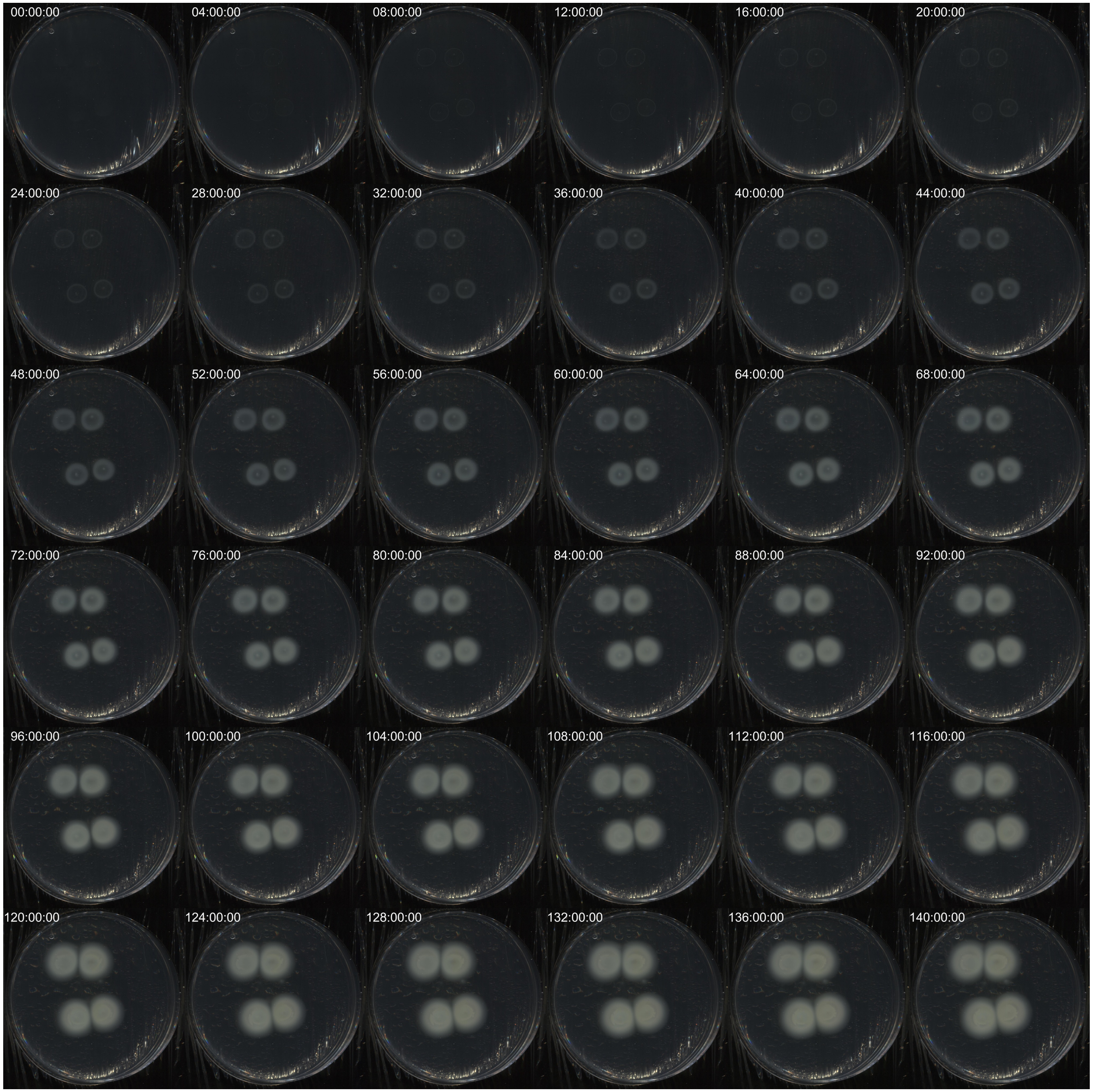


Figure S2 Four colony setup of *B. diazoefficiens* USDA110 in soft agar, imaged every 4h from d0 to d6

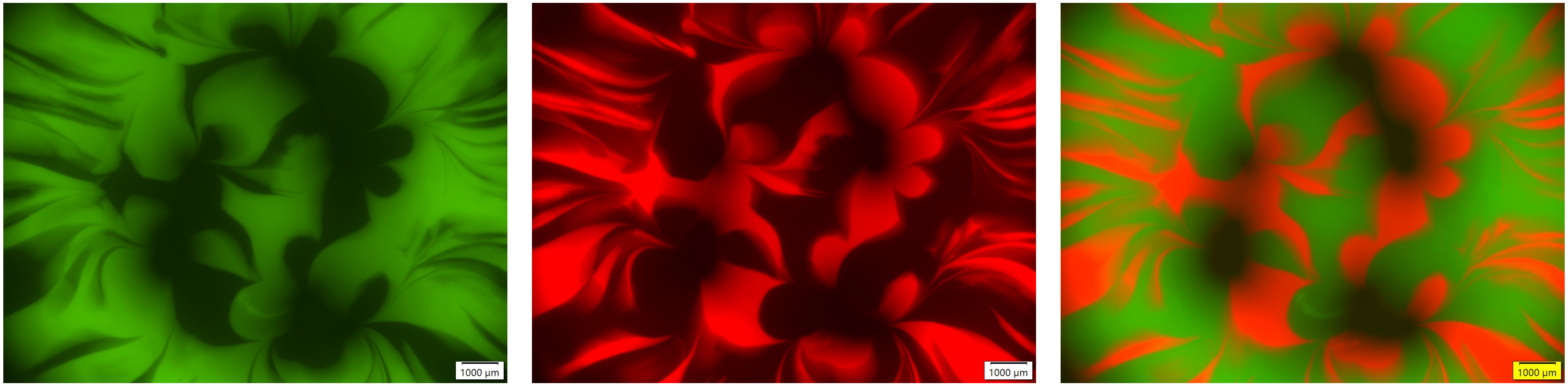


Figure S3 Fluorescence micrograph of region of a colony of bj-GFP and mCherry-tagged strains of *B. diazoefficiencs* USDA110 grown on 1.5% agar.

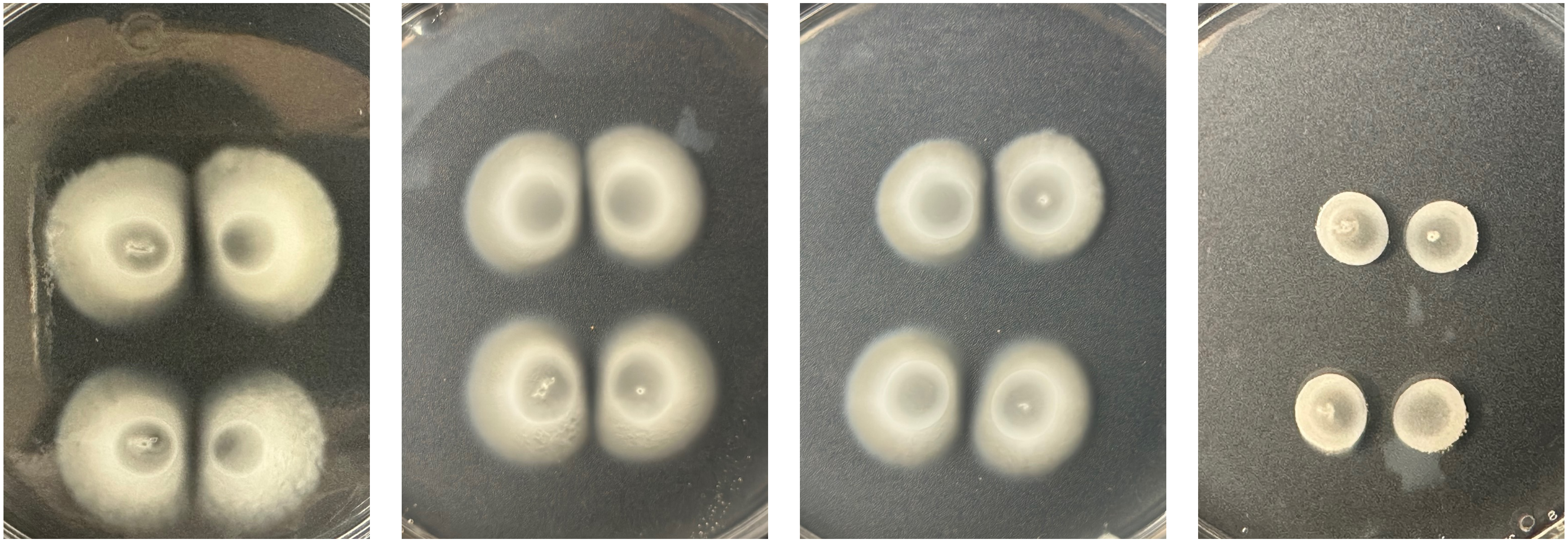

**Wild type** | **Δ*fliC*** | **Δ*lafA*** | **Δ*fliC-lafA***

Figure S4 Four colony set-up of *Bradyrhizobium* USDA110-Spc4, and its *fliC, lafA* and *fliC-lafA* deletion mutants, incubated in PSY soft agar with arabinose for 7 d.

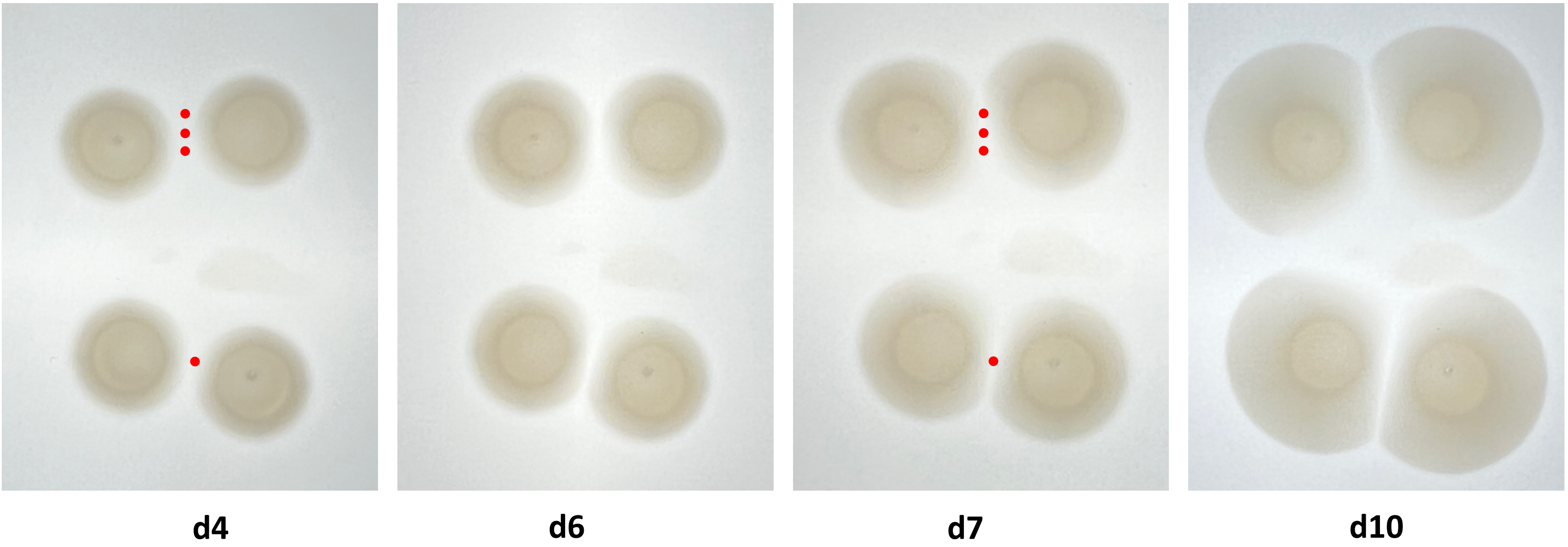


Figure S5  Four colony set-up of *Bradyrhizobium* USDA110 in PSY soft agar with arabinose, with 5µL of 10X PSY added on d 4 and again d7 at the positions marked by red dots.

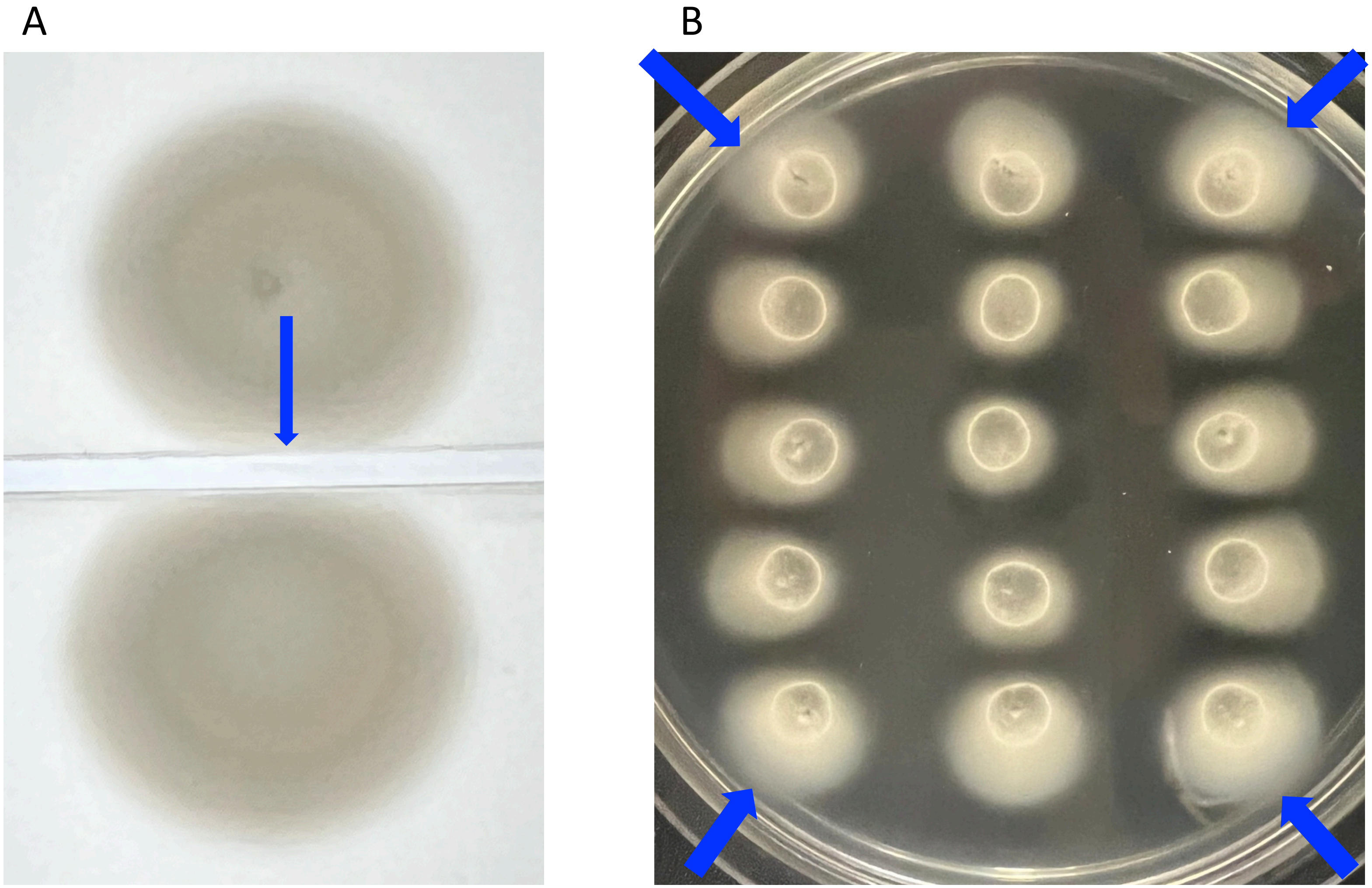


Figure S6  Swimming colonies of *Bradyrhizobium* USDA110 in PSY soft agar with arabinose, with a sterile glass slide placed between them (a), and colonies spotted close to the edge of a Petri dish (b).
